# Circadian clock factors regulate seed oil accumulation by promoting fatty acid synthesis in Arabidopsis

**DOI:** 10.1101/2022.12.15.520653

**Authors:** Sang-Chul Kim, Kristen N. Edgeworth, Dmitri A. Nusinow, Xuemin Wang

## Abstract

The circadian clock regulates temporal metabolic activities, enabling organisms to adapt to cyclic environmental changes, but how it affects lipid metabolism in plants is poorly understood. Our previous finding showed that the central clock transcription factors LATE ELONGATED HYPOCOTYL (LHY) and CIRCADIAN CLOCK ASSOCIATED 1 (CCA1) increased seed oil contents in Arabidopsis. Here we investigated the molecular and metabolic mechanism underlying the LHY and CCA1 regulated oil accumulation. Triacylglycerol (TAG) accumulation in Arabidopsis developing seeds was increased in *LHY*-overexpressing (LHY-OE) and decreased in *lhycca1* plants compared to wild-type (WT). Metabolic tracking of lipids in developing seeds indicated that fatty acids (FAs) of major lipid precursors for TAG production increased more rapidly in LHY-OE and slowly in *lhycca1* than in WT, suggesting that LHY enhanced FA synthesis. Transcript analysis revealed that the expression of genes involved in FA synthesis, including the one encoding β-ketoacyl-ACP synthase III (KASIII), was oppositely changed in developing seeds of LHY/CCA1-OEs and those of *lhycca1*. Chromatin immunoprecipitation, electrophoretic mobility shift, and transactivation assays indicated that LHY directly bound and activated the promoter of *KASIII*. Furthermore, phosphatidic acid, a metabolic precursor to TAG, inhibited LHY binding to *KASIII* promoter elements. Our data reveal a new regulatory mechanism by the core clock regulators for storage lipid production during plant seed development.

## INTRODUCTION

The circadian clock is a self-sustaining endogenous oscillator that orchestrates 24-hour rhythms of cellular metabolism, organismal physiology, and behavior, enabling plants to anticipate and adapt to daily environmental cycles. The molecular oscillator consists of multiple interlocking transcriptional and posttranslational feedback loops composed of transcription factors that regulate the genes of one another reciprocally and downstream target genes mediating numerous biological processes (Hsu and Harmer, 2014; McClung, 2019). The central loop of the molecular clock in Arabidopsis (*Arabidopsis thaliana*) is orchestrated by LATE ELONGATED HYPOCOTYL (LHY) and its closely related and functionally redundant CIRCADIAN CLOCK ASSOCIATED1 (CCA1) and their target TIMING OF CAB EXPRESSION1 (TOC1) (McClung, 2019). LHY and CCA1 accumulate in the morning and suppress *TOC1* expression by binding to its promoter, while their expression is suppressed reciprocally by TOC1 accumulation in the evening (Alabadí et al., 2001; Gendron et al., 2012; Huang et al., 2012). LHY/CCA1 and TOC1 function as transcriptional repressors not only for each other but also for various target genes, including central clock genes, such as *PSEUDO-RESPONSE REGULATOR*s (*PRR*s), *GIGANTEA* (*GI*), *LUX ARRHYTHMO* (*LUX*), and *EARLY FLOWERING 3 and 4* (*ELF3* and *ELF4*) (Nusinow et al., 2011; Huang et al., 2012; Herrero et al., 2012; Kamioka et al., 2016), and multiple output genes including those for flowering time regulation, hypocotyl elongation, and cold stress responses (Niwa et al., 2009; Nakamura et al., 2014a; Kidokoro et al., 2021). Thus, the circadian clock is a highly complex network of the Arabidopsis clock regulatory circuit consisting of a remarkably large number of molecular components, mainly transcription factors and their co-regulators (McClung, 2019; Nohales, 2021).

Among a wide range of metabolic effects of the circadian clock, lipid metabolism is of particular interest, especially in the medical context of human research. Temporal stress caused by clock misalignments, such as shift work, chronic jet lag, and sleep deprivation, and clock gene mutations are associated with various lipid metabolic diseases, such as obesity and non-alcoholic fatty liver disease (Shi et al., 2013; Panda, 2016; Chaix et al., 2019). In plants, the circadian regulation of lipid content and composition is an emerging issue, and its functional and physiological significance is in the process of being unraveled. It is a general observation that contents of polyunsaturated fatty acids (FAs), such as linoleic acid (18:2) and linolenic acid (18:3), in total lipids and membrane glycerolipids (particularly phosphatidylcholine; PC) increase during the night and decrease during the day at the expense of oleic acid (18:1), as demonstrated in many plant species, including cotton, spinach, and Arabidopsis (Browse et al., 1981; Rikin et al., 1993; Ekman et al., 2007; Maatta et al., 2012; Nakamura et al., 2014b). In addition, Arabidopsis lipidomic analyses revealed that membrane lipid classes phosphatidylserine (PS), phosphatidic acid (PA), and chloroplast galactolipids accumulated at nighttime (Maatta et al., 2012), and consistently that the levels of some molecular species of PS, PA and chloroplast-enriched phosphatidylglycerol (PG) oscillated with a period of ∼24 hours in wild-type (WT) plants, but not in *lhycca1* (Kim et al., 2019). The levels of PC species showed clear daily changes, with those containing less unsaturated FAs (34:1 and 36:1∼4) predominant in the daytime and 18:3-containing species (36:5 and 36:6) increased during the night (Nakamura et al., 2014b). Interestingly, a component of florigen FLOWERING LOCUS T (FT) bound preferentially to the day-dominant PC species to promote flowering, indicating a physiological effect of daily lipid changes (Nakamura et al., 2014b). It was also reported that some lipid metabolic genes in Arabidopsis were under circadian control and expressed differently in *lhycca1* and *CCA1*-overexpressing plants (CCA1-OE; Hsiao et al., 2014; Nakamura et al., 2014a; Nagel et al., 2015). However, it is unclear whether and how this differential gene expression leads to circadian changes in the lipid accumulation mentioned above.

As such, the circadian regulation of plant lipid metabolism has been studied mainly in the context of polar membrane glycerolipids in vegetative tissues. However, we recently found that the accumulation of Arabidopsis seed oil decreased in *lhycca1* and increased in *LHY*-overexpressing plants (LHY-OE) compared to WT. In contrast, no significant difference was observed in *lhy*, possibly due to the overlapping function of CCA1 (Kim et al., 2019). Similarly, CCA1-OE seeds also showed a substantial oil content increase (Hsiao et al., 2014). Those results suggest the circadian regulation of storage lipids during seed development in addition to membrane lipids during vegetative growth. The primary storage oil in seeds or fruits is triacylglycerol (TAG), composed of a glycerol backbone with three esterified FAs. TAG is stored as a high-energy compound and broken down to provide energy for the growth of young seedlings during the early stage of seed germination until they begin to photosynthesize energy compounds (Yang and Benning, 2018). In addition to being essential to plant growth, vegetable oils are an important source of energy for human and animal diets and provide feedstocks for biofuels and industrial products. Thus, a better understanding of how circadian clock regulators influence storage lipid metabolism has the potential to enhance the understanding of how the clock regulates storage lipid synthesis and accumulation and improve the production of plant oils for numerous applications. Here, we investigated how the central clock transcription factors influence TAG accumulation in developing seeds of Arabidopsis.

## RESULTS

### LHY and CCA1 affect Arabidopsis seed oil content and composition

To investigate the effects of LHY and CCA1 on seed oil accumulation, we used double knockout mutant *lhycca1* and CaMV-35S promoter-driven *LHY*- and *CCA1*-OE. We first confirmed the plant materials by using molecular and phenotypic analyses. Reverse transcription (RT)-PCR and quantitative real-time (qRT)-PCR analyses revealed that no transcript of *LHY* and *CCA1* was detected in two independent plants of *lhycca1* (Figure S1A). In WT plants, both genes were highly expressed at Zeitgeber time 0 hour (ZT0) and expressed at a much lower level at ZT12 (Figure S1A), consistent with their previously documented rhythmic expression patterns (Alabadí et al., 2001; Gendron et al., 2012; Huang et al., 2012). Immunoblotting of total nuclear proteins isolated from *LHY/CCA1*-OEs demonstrated ZT-independent enrichment of LHY and CCA1 proteins, verifying the constitutive overexpression of *LHY* and *CCA1* in the transgenic plants (Figure S1B). Clock perturbation by the overexpression of *LHY* or *CCA1* has been reported to result in abnormal phenotypes, including long hypocotyls and late flowering, that are highly distinguishable from those of WT (Mizoguchi et al., 2002; Niwa et al., 2009; Lu et al., 2012). We indeed observed the characteristic morphological and developmental phenotypes in *LHY/CCA1*-OEs; when compared to WT, they had approximately three times longer hypocotyls (Figure S1C) and began to flower about two weeks later, while *lhycca1* exhibited ∼10-day earlier emergence of the first flower (Figure S1D). Our analyses of the circadian leaf movement revealed that the period was shortened in *lhycca1* (∼18 hours), and *LHY/CCA1*-OEs were arrhythmic, whereas WT plants displayed the typical 24-hour oscillation (Figure S1E). The data indicate that they are highly consistent with previous studies and that the plant materials are authentic clock-perturbed lines.

Gas chromatography-based measurements of the *LHY*/*CCA1*-altered seeds revealed that oil content was decreased by approximately 15% in *lhycca1* and increased by up to 25% in *LHY/CCA1*-OEs (Figure 1A). The differential accumulation of seed oil in *LHY*/*CCA1*-altered plants was also demonstrated by our analyses using nuclear magnetic resonance (Figure S1F). Compared with WT, *LHY*-or *CCA1*-OE seeds displayed an increase in 18:1, 18:2, 18:3, and eicosenoic acid (20:1), whereas *lhycca1* seeds had a decrease in 18:1, 18:2, and 20:1, but not 18:3 (Figure 1B).

**Figure 1.**
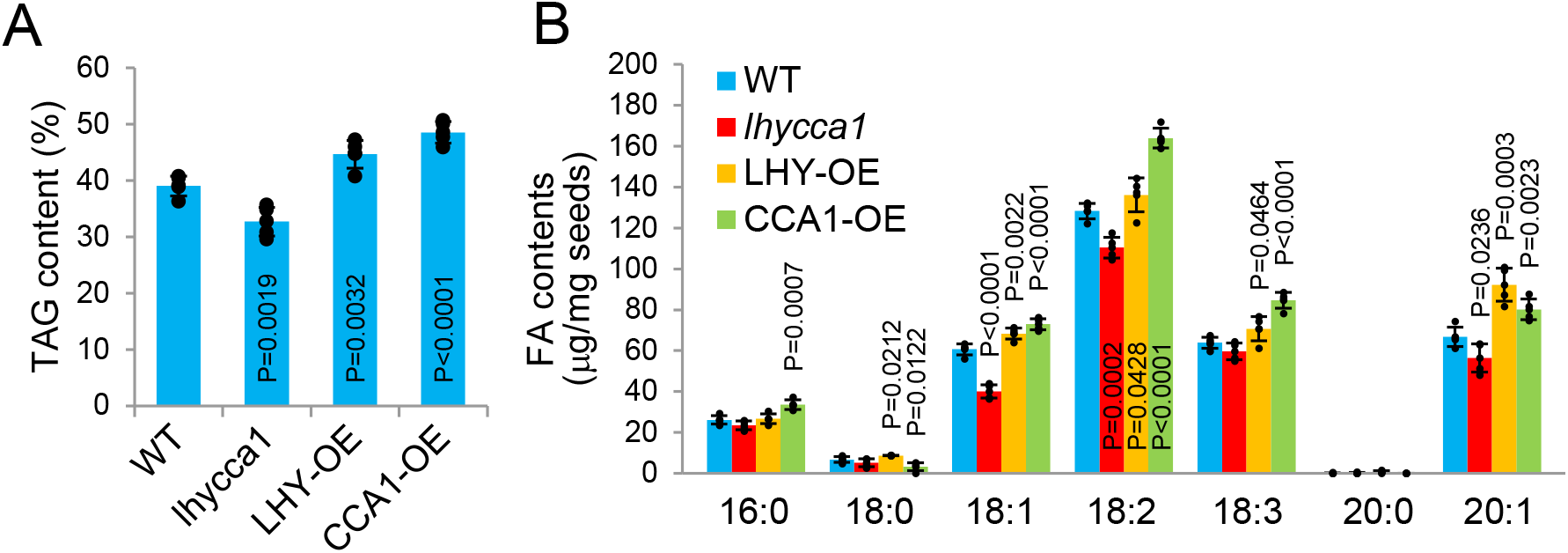
Seed oil content in *LHY*/*CCA1*-altered plants. **A**. Total TAG content of dry seeds from the indicated plant lines measured by gas chromatography as expressed as % (w/w) of total seed weight. **B**. Contents of individual fatty acids. The data in A are shown here as the amounts of individual species of fatty acids (number of carbons : number of double bonds). Values in both panels are mean ± SD with individual data points and *p* values denoting statistical significance (to WT; n=5).

### LHY/CCA1 increases TAG accumulation during seed development

To assess LHY and CCA1’s effects on TAG accumulation during seed development, we measured TAG content over time in developing seeds of *lhycca1* and *LHY*-OE plants from 5 days after flowering (DAF) up to 13 DAF. TAG content began to increase after 7 DAF and reached the maximum at ∼11 DAF in WT and *LHY*-OE, whereas the peak in *lhycca1* was significantly delayed (Figure 2AB). As a result, TAG was accumulated more rapidly in *LHY*-OE but slower in *lhycca1* than in WT during the stage of the most active TAG formation (9∼11 DAF), as evidenced by the statistical significance in the slopes of lines between the two consecutive time points (Figure 2B). During this stage, the accumulation rate of 18:0-containing TAG was decreased in *lhycca1*, whereas that for 18:1 or 18:2 TAG was increased in *LHY*-OE, compared with WT developing seeds (Figure 2C). The data indicate that *LHY* and *CCA1* increase TAG accumulation at the early seed maturation stage.

**Figure 2.**
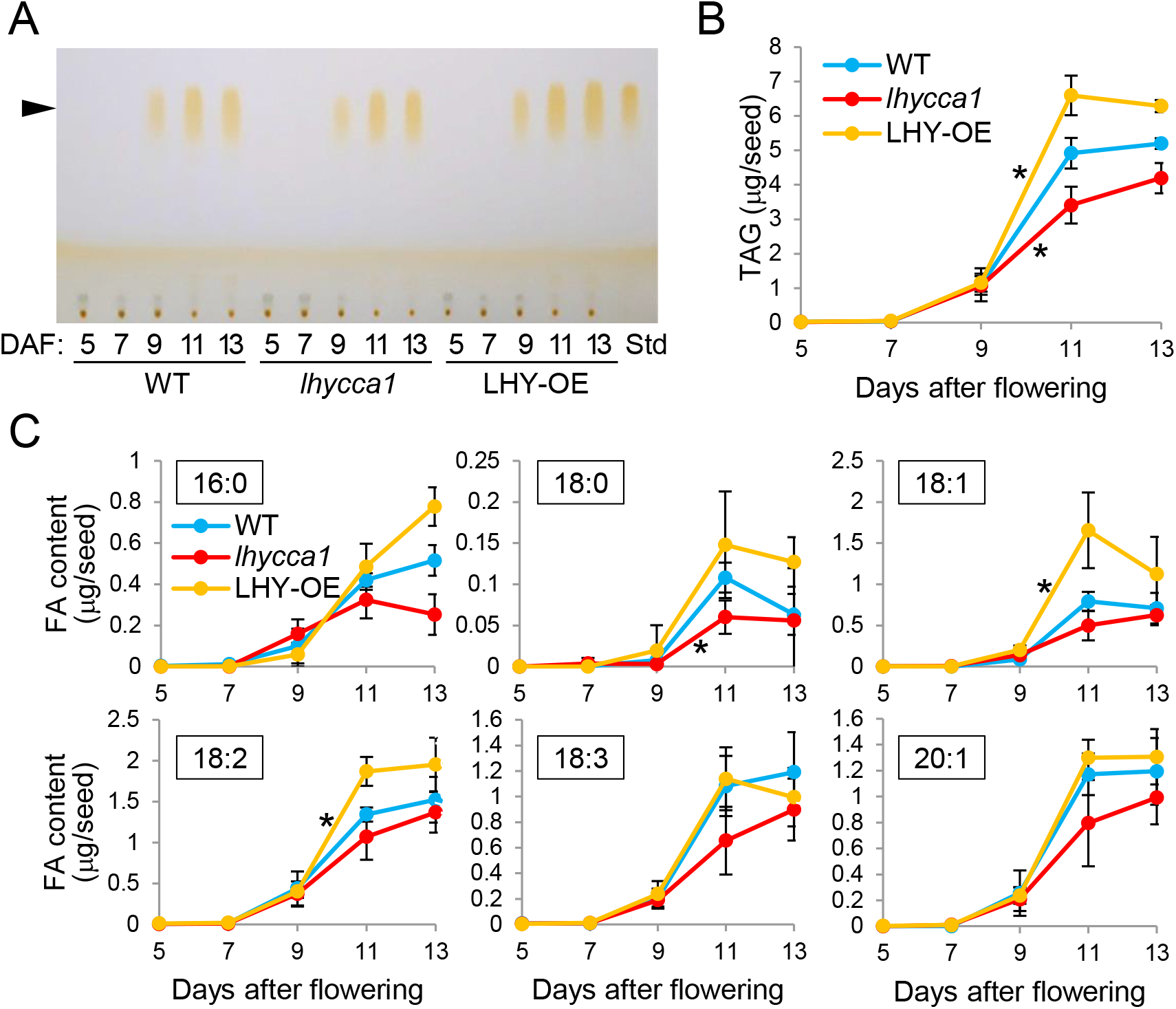
TAG accumulation during seed development. **A**. Thin layer chromatography (TLC) separation of TAG. Total lipids were extracted from developing seeds of WT, *lhycca1*, and LHY-OE at the indicated days after flowering (DAF). The lipids were separated by TLC and visualized by iodine exposure. Arrowhead indicates the position of TAG. Std, authentic TAG standard. **B**. Contents of total TAG. TAG spots in A were scraped out and quantified by gas chromatography for TAG contents. Values are mean ± SD (n=3). Asterisk denotes statistical significance of the slope compared to that of WT (*p*<0.05). **C**. Contents of individual fatty acids. The data in B are shown here as the amounts of individual species of fatty acids (number of carbons : number of double bonds). Values are mean ± SD (n=3).

In addition, we measured changes in TAG contents over time during seed germination up to 5 days after imbibition (DAI). TAG began to be broken down at 1 DAI and was almost completely depleted at 4 DAI in all plant lines tested (Figure S2AB). Compared to WT, there was no significant difference in TAG degradation rates in *lhycca1* or *LHY*-OE, including the stage of the most robust degradation (1∼2 DAI; Figure S2B). Individual FAs displayed highly similar patterns of progressive decline in their levels among *lhycca1*, LHY-OE, and WT (Figure S2C). These data suggest that the alterations of *LHY* and *CCA1* expression have no apparent effect on TAG degradation in germinating seeds.

### TAG metabolic tracking indicates LHY/CCA1 promotion of FA synthesis

To determine how *LHY*- and *CCA1*-alterations affect TAG content, we used [^14^C]-acetate to track the production of TAG and its immediate precursors diacylglycerol (DAG) and PC in developing seeds from WT, *lhycca1*, and LHY-OE plants. Developing 10-DAF seeds were used because they were most active in TAG production (Figure 2B). [^14^C]-Acetate is converted to [^14^C]-acetyl-CoA and labels newly synthesized FAs that are used to produce TAG, using the Kennedy pathway involving sequential acylation of a glycerol backbone and/or other reactions involving PC metabolism (Figure 3A). First, we performed ‘pulse labeling’ to monitor the production of lipids, where their radioactivity was measured over time during the incubation of seeds with the radioisotope. The radioactivity of all three lipids increased continuously without saturation of the lipid pools (Figure 3B). When the three lipids were compared for each of the plants, the incorporation rate of [^14^C]-acetate was always in the order of TAG, PC, then DAG, which was observed consistently in WT, *lhycca1*, and *LHY*-OE seeds (Figure S3). TAG, DAG, and PC were labeled with [^14^C]-acetate more rapidly in *LHY*-OE and slowly in *lhycca1* than in WT (Figure 3B).

**Figure 3.**
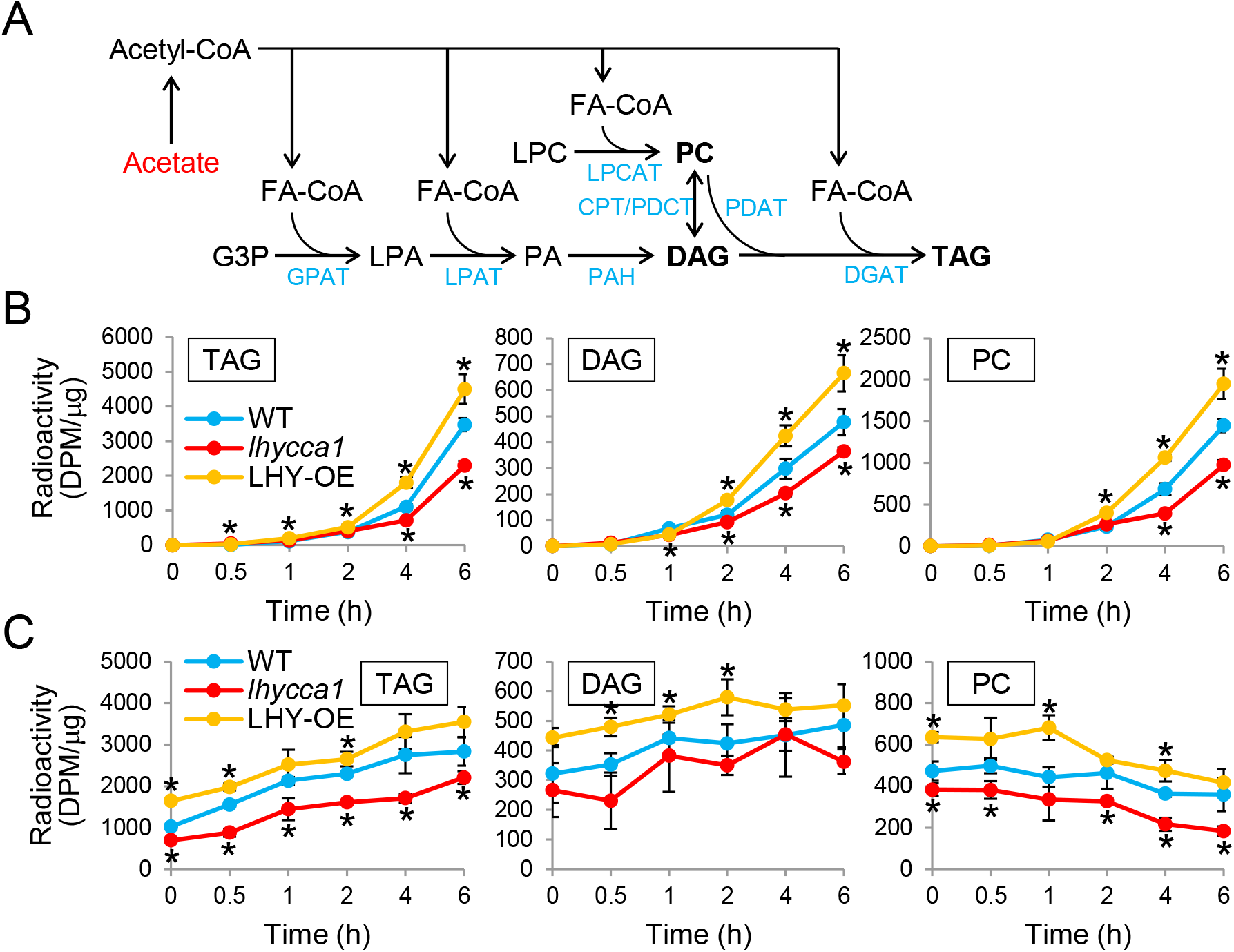
Metabolic tracking of radio-labeled lipid precursors of TAG. **A**. TAG biosynthetic pathways. Radioactive substrate used in this study is highlighted in red. Enzymes are in blue and the major lipid precursors (PC and DAG) and the final product (TAG) are in bold. G3P, glycerol-3-phosphate; LPA, lysoPA; LPC, lysoPC; GPAT, G3P acyltransferase; LPAT, LPA acyltransferase; DGAT, acyl-CoA:DAG acyltransferase; PDAT, PC:DAG acyltransferase; PAH, PA phosphohydrolase; LPCAT, LPC acyltransferase; CPT, choline phosphotransferase; PDCT, PC:DAG Cholinephosphotransferase. **B**. Pulse labeling. 10-day after flowering (DAF) seeds of WT, *lhycca1*, and LHY-OE were incubated with [^14^C]-acetate for the indicated time. Total lipids were extracted and separated by thin layer chromatography. Radioactivities of TAG, DAG, and PC were measured and are shown here as per μg chlorophyll of the seeds. Values are mean ± SD (n=5). Asterisk denotes statistical significance compared to WT (*p*<0.05). **C**. Pulse-chase labeling. 10-DAF seeds pre-incubated with [^14^C]-acetate for 3 h were washed to remove radioisotopes and further incubated for the indicated time. Lipids were processed and analyzed as in B. Values are mean ± SD (n =5).

In addition, we performed ‘pulse-chase labeling’ to monitor lipid turnover. Developing seeds were first labeled with [^14^C]-acetate, then washed to remove the labeling radioisotope, and further incubated without the radioisotope for time-course measurement of lipid radioactivity. For TAG, DAG, and PC, the differential FA labeling (*LHY*-OE > WT > *lhycca1*) observed in the pulse labeling remained essentially unchanged during the chase for up to 6 hours (Figures 3C and S4). This suggests that the lipids were turned over at a similar rate in WT, *lhycca1*, and LHY-OE, and further details on the labeling data interpretation will follow in the ‘Discussion’ section. Our data suggest that LHY and CCA1 increase TAG accumulation by promoting FA synthesis in developing seeds rather than suppressing its degradation.

### LHY/CCA1 affects the expression of genes for FA biosynthesis

Based on our data indicating that *LHY*/CCA1-OE enhances FA production, we measured the expression levels of the genes encoding some key FA synthetic enzymes in 10-DAF seeds of the *LHY*/*CCA1*-altered plants. The first committed step of FA synthesis is catalyzed by acetyl-CoA carboxylase (ACC), which converts acetyl-CoA to malonyl-CoA (Figure 4A). Arabidopsis has two types of ACCs: the cytosolic eukaryotic ACCs (ACC1 and ACC2) involved in FA elongation and the plastidic prokaryotic ACC responsible for *de novo* FA synthesis. The plastidic ACC is composed of multiple subunits: biotin carboxylase (BC), biotin carboxyl carrier protein (BCCP), and α- and β-carboxyltransferase (CT) (Figure 4B). In addition, there present biotin attachment domain-containing (BADC) proteins that compete with BCCP and inhibit ACC activity (Figure 4B; Salie et al., 2016; Ye et al., 2020). The transcript level of most subunits of the plastidic ACC was largely similar in the developing seeds of WT, *lhycca1*, and LHY/CCA1-OE (Figure 4C). However, the transcript level of *BCCP2* and *BADC2* showed an opposite change in *lhycca1* and LHY/CCA1-OE. *BCCP2* expression was reduced in *lhycca1* and increased in LHY/CCA1-OEs and conversely, *BADC2* exhibited highly increased and decreased expression in *lhycca1* and LHY-OE, respectively (Figure 4C). By comparison, the transcript level of cytosolic *ACCs* was comparable in the developing seeds of *LHY*- and/or *CCA1*-OE, *lhycca1*, and WT (Figure 4D).

**Figure 4.**
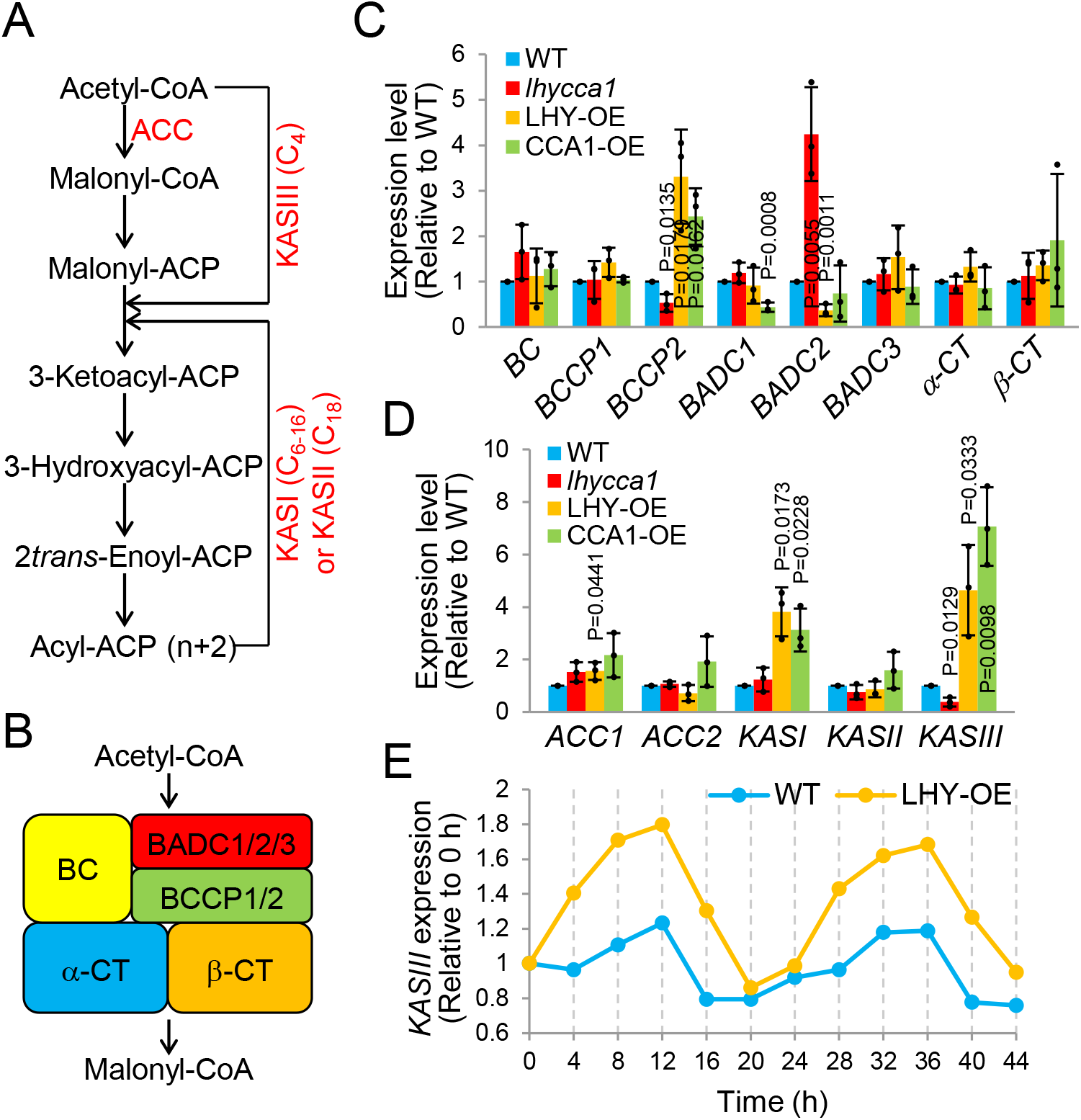
Expression of the genes required for fatty acid synthesis. **A**. Fatty acid biosynthetic pathway. Intermediate enzymes, cofactors, and byproducts are omitted for simplicity. The chain length (number of carbons) of fatty acids produced by each KAS is in parenthesis. ACC, acetyl-CoA carboxylase; ACP, acyl carrier protein. **B**. Subunits of heteromeric ACC. Note that relative size and position of the subunits are different from reality. See text for abbreviations. Numbers refer to isoforms in Arabidopsis. **C**. Expression levels of plastidic *ACC* subunits. Total RNA was extracted from 10-DAF seeds of WT, *lhycca1*, and LHY/CCA1-OEs. Quantitative real-time PCR was performed with gene-specific primers. Values are mean ± SD with individual data points and *p* values denoting statistical significance (to WT; n=3). **D**. Expression levels of cytosolic *ACC* and *KAS*. The experiments performed and data layout are as in C. **E**. Time-course expression of *KASIII*. The data were generated from a public microarray database (‘Diurnal’; diurnal.mocklerlab.org), where WT (*Ler*) and LHY-OE were grown under 8-h light/16-h dark cycles and are shown here as normalized to the level at 0 h. Pearson’s correlation coefficient: WT=0.872, LHY-OE=0.979.

The malonyl group is condensed with acetyl-CoA by β-ketoacyl-ACP synthase (KAS) III to produce a 4-carbon FA, whereas KASI uses the 4-carbon FA and malonyl groups to produce up to 16-carbon FAs, and KASII elongates 16-carbon acyl-ACP to produce 18-carbon FAs (Figure 4A). *KASII* transcript level was similar in *LHY/CCA1*-OE, *lhycca1*, and WT, whereas *KASI* was increased in *LHY/CCA1*-OE but had no difference in *lhycca1* compared to WT (Figure 4D). However, the expression of *KASIII* displayed opposite changes in *lhycca1* and *LHY/CCA1*-OE. *KASIII* transcript level was approximately 5-to 8-fold higher in *LHY/CCA1*-OE, but 3-fold lower in *lhycca1* than in WT (Figure 4D). Analysis of a microarray gene expression database (diurnal.mocklerlab.org) indicates that *KASIII* displayed rhythmic expression with a period of ∼24 hours, and its magnitude was increased considerably in *LHY*-OE (Figure 4E). The altered patterns of expression suggest that the early steps of FA synthesis mediated by KASIII and/or BCCP2/BADC2 may be subject to the LHY/CCA1 regulation.

### LHY binds the KASIII promoter and enhances its expression

Based on the gene expression data, we analyzed whether the genomic sequences of *KASIII, BCCP2*, and *BADC2* contain potential binding sites of LHY/CCA1. The promoter region of *KASIII*, but not *BCCP2* or *BADC2*, has two evening elements (EEs) and CCA1-binding sites (CBSs) (Figure 5A). EE and CBS are two *cis*-elements bound by both LHY and CCA1, which have been found in many genes regulated by these transcription factors (Green and Tobin, 1999; Farré et al., 2005; Nagel et al., 2015; Kamioka et al., 2016). Thus, we carried out chromatin immunoprecipitation (ChIP) to isolate CCA1 from *CCA1*-OE followed by PCR to amplify co-precipitated DNA. *KASIII* promoter was effectively amplified with two of three primer pairs (P1 and P3) from ChIP DNA from two independent *CCA1*-OE plants, but not from WT, consistent with the binding observed for the *TOC1* promoter (Figure 5B). As expected, *BCCP2* and *BADC2* were not clearly detected with any of four primer pairs binding across their promoters (Figure 5B).

**Figure 5.**
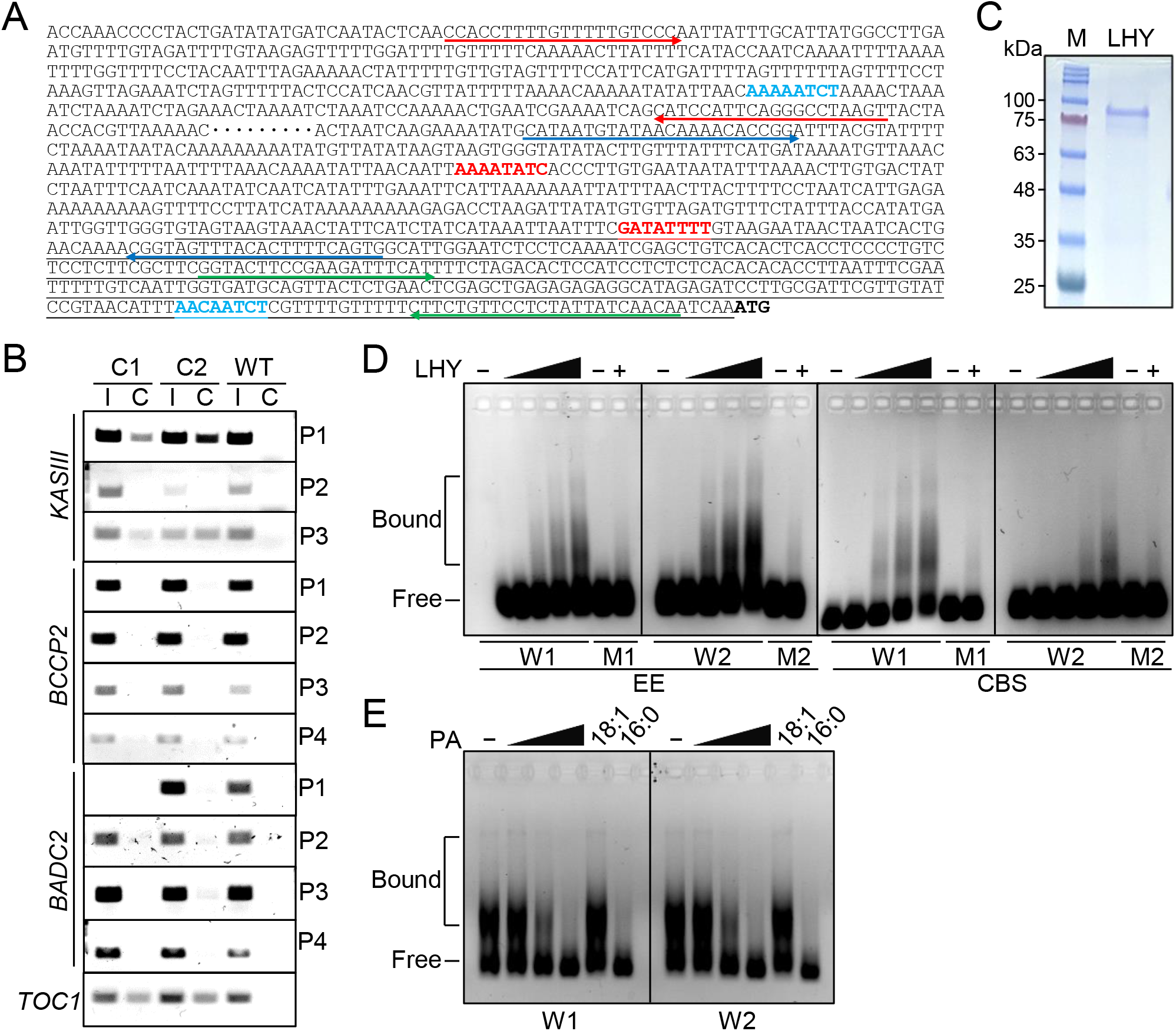
LHY/CCA1 binding to *KASIII* promoter. **A**. DNA sequence of *KASIII* promoter region. 1,469 bps upstream from transcription start site (1,825 bps from start codon) is shown with omission of some intervening nucleotides (….). 5’-UTR is underlined and the start codon is in bold. EE and CBS are in red and blue, respectively. Arrows indicate primer binding sites for ChIP-PCR (red, P1; blue, P2; green, P3). **B**. ChIP-PCR results. DNA immunoprecipitated with CCA1 was amplified by PCR using primers indicated on the right for the gene promoters on the left. *TOC1* at the bottom is a positive control. C1 and C2, two independent *CCA1*-OEs; I, input DNA; C, ChIP DNA. **C**. Purified LHY protein. LHY was expressed in and purified from *E. coli* and subjected to SDS-PAGE followed by Coomassie blue staining. Size of marker proteins (M) is on the left. **D**. Electrophoretic mobility shift assay. 60-bp DNA fragments containing wild-type (W) or mutated (M) EE and CBS were incubated with the purified LHY and separated in agarose gel. Triangle indicates an increasing amount of LHY (1∼20 μg). − and + denote 0 μg and 20 μg, respectively. **E**. PA effect on LHY binding to EE. EMSA was performed as in C with wild-types EEs and LHY (20 μg). − denotes no PA. Triangle indicates an increasing amount of PA mixture (from egg yolk; 0.1∼10 μg). 18:1 and 16:0 indicate the acyl chain at *sn*-1 position of PA species with 18:1 at *sn*-2.

We also performed an electrophoretic mobility shift assay (EMSA) to test the *in vitro* binding of LHY to EE and CBS binding sites in the *KASIII* promoter. LHY protein was expressed and purified from *Escherichia coli* to near homogeneity (Figure 5C). For both EE and CBS, the protein bound effectively to ∼60-bp DNA fragments containing each of the two sites (W1 and W2), but not of those mutated (M1 and M2) in an LHY dose-dependent manner, as indicated by the slower electrophoretic migration of a portion of the DNA (Figure 5D). A control protein, glyceraldehyde-3-phosphate dehydrogenase, did not affect the electrophoretic mobility of the same DNA fragments (Figure S5). In addition, based on our previous finding that specific PA species bound to and inhibited LHY/CCA1 interaction with their target promoters (Kim et al., 2019), we examined the PA effect on LHY binding to the two EEs in *KASIII* promoter by EMSA. The binding was severely inhibited by mixed PA species from egg yolk in a dose-dependent manner. We showed previously that 16:0-18:1 PA bound to LHY, whereas 18:1-18:1 PA did not (Kim et al., 2019). The 16:0-18:1 PA inhibited the LHY binding to *KASIII* promoter elements, but the non-binding 18:1-18:1 PA had no effect on the binding (Figure 5E), suggesting that PA-LHY binding is required for the PA inhibition of LHY-*KASIII* promoter interaction.

We then investigated the effect of LHY on the *KASIII* promoter by performing transactivation assays, where the *KASIII* promoter or *TOC1* promoter drove the expression of luciferase (LUC) reporter as a control (Figure 6A). The effector LHY was expressed by the CaMV-35S promoter as a fusion with green fluorescence protein (GFP) that was used as an internal control for normalization (Figure 6A). All possible combinations of the effectors and reporters were co-expressed transiently in tobacco leaves, and fluorescence (GFP) and luminescence (LUC) signals of their protein extracts were recorded. The successful transformation of all samples was confirmed by PCR amplification of the respective promoters and by the GFP intensity being significantly greater than that of a non-transformant control (Figure 6B). Compared to controls, an intense LUC signal was detected from the sample transformed with the test effector and test reporter (T_E_T_R_; Figure 6B), which was also observed when the signal intensity was background-subtracted and normalized to the GFP signal (Figure 6B). It is worth noting that T_E_C_R_ lowers the LUC signal compared to C_E_C_R_, as expected, as LHY functions as a repressor when bound to the *TOC1* promoter (Figure 6B). The data here indicate that the *KASIII* promoter is directly associated with and activated by LHY in plants.

**Figure 6.**
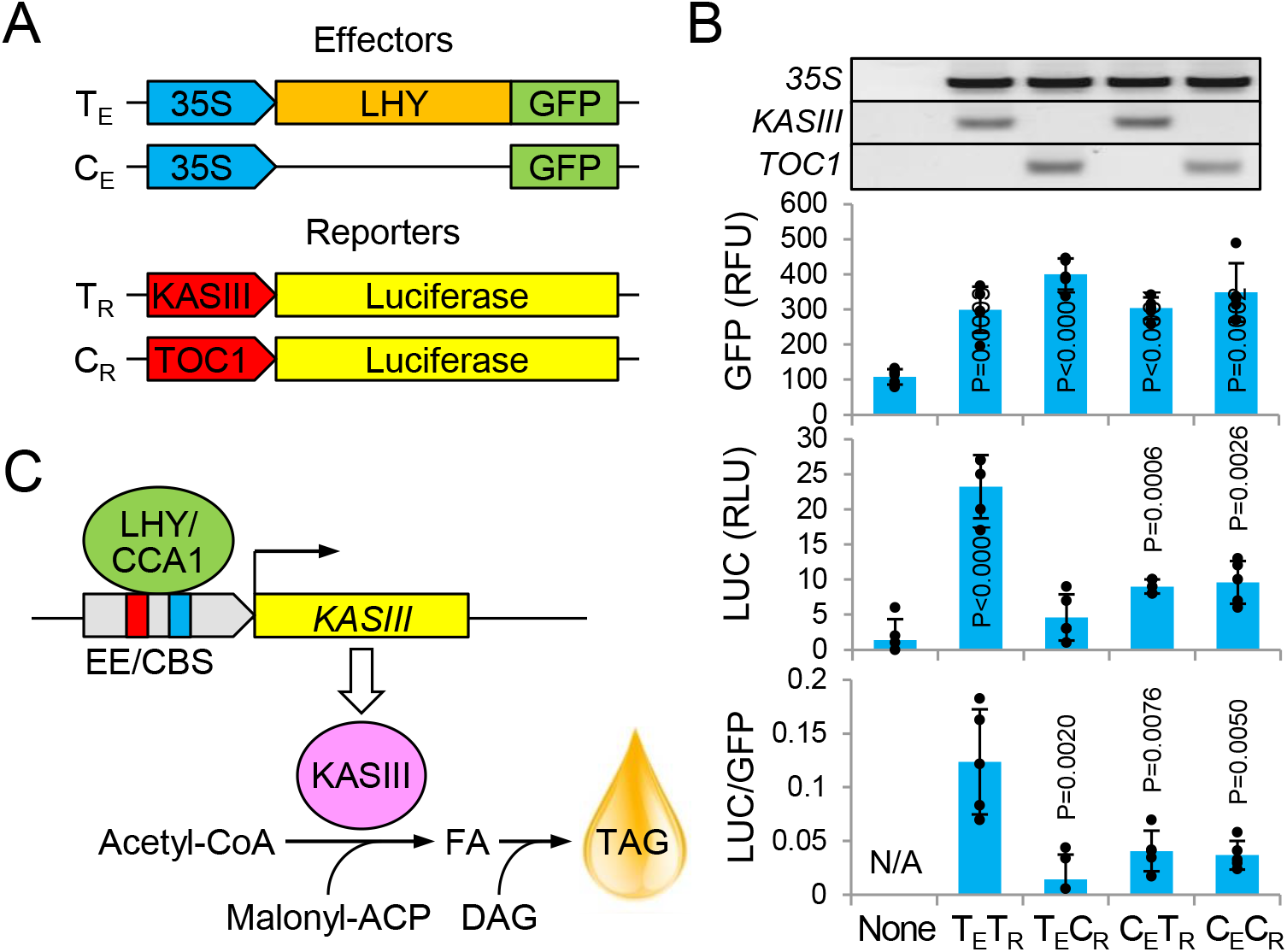
LHY/CCA1 activation of *KASIII* promoter. **A**. Schematic diagram of the reporter assay vectors. Effector (E) and reporter (R) genes and their promoters are shown. T, test vector; C, control vector. **B**. Fluorescence and luminescence intensities. Tobacco leaves were transformed with the indicated combinations of vectors. Gel images are PCR results from the transformed tobacco DNA with the primers indicated on the right. Total proteins extracted were measured for intensities of fluorescence (GFP) and luminescence (LUC). At the bottom is baseline (None)-subtracted values shown as LUC normalized to GFP. Values are mean ± SD with individual data points and *p* values denoting statistical significance (to None for GFP and LUC; to T_E_T_R_ for LUC/GFP; n=5). RFU, relative fluorescence unit; RLU, relative luminescence unit; None, non-transformant; N/A, not applicable. **C**. Proposed model for LHY/CCA1-regulated seed oil accumulation. LHY and CCA1 enhance *KASIII* expression by binding to *KASIII* promoter through its EE and/or CBS. KASIII then catalyzes the synthesis of FAs that are incorporated into TAG during seed development. EE, evening element; CBS, CCA1-binding site.

## DISCUSSION

The circadian clock regulates various metabolic and physiological processes, and its disruption in mammals leads to metabolic disorders such as increased lipid accumulation and diabetes. In plants, the clock-disrupted mutants *lhycca1* and LHY/CCA1-OEs altered storage lipid production in Arabidopsis seeds. However, how the perturbations of the clock factors lead to altered lipid accumulation remained unknown. This study identifies a mechanism by which LHY/CCA1 regulates storage lipid production. Our data indicate that those key clock regulators bind to the *KASIII* promoter to enhance the expression of *KASIII* that mediates the initial condensation reaction of FA synthesis and promotes FA synthesis for TAG production during seed development (Figure 6C).

Our data from metabolic analyses provide further support for LHY/CCA1’s promotion of FA synthesis. When labeling rates of TAG, DAG, and PC were compared side by side for WT, *lhycca1*, and *LHY*-OE, TAG was labeled with [^14^C]-acetate most rapidly during the entire pulse period. These labeling patterns are consistent with the current model of precursor-product relationships in the Kennedy pathway (Figure 3A) and with previous data from various plant species, including Arabidopsis (Lu et al., 2009; Bates and Browse, 2011; Bates et al., 2012; Bates et al., 2014; Allen et al., 2015; Yang et al., 2017), indicative of the experimental validity of our analyses. Such typical labeling patterns were observed consistently in all three plant lines, suggesting that LHY/CCA1 had no apparent effect on the precursor-product relationships of TAG, DAG, and PC (Figure S3). Comparing the three plants for each of the three lipids, on the other hand, the accumulation rates of FA-labeled TAG, DAG, and PC in the pulse labeling were increased in *LHY*-OE and decreased in *lhycca1* (hereafter designated as OE and KO, respectively). After being synthesized in plastids and bound to CoA in the cytosol, FAs are incorporated into the three glycerolipids by distinct enzymatic routes in the endoplasmic reticulum (ER) (Figure 3A). The same FA labeling pattern (OE>WT>KO) of all three lipids, despite their different FA incorporation routes, indicates that LHY/CCA1 may enhance FA synthesis rather than any steps occurring in the ER. If the clock proteins specifically increased the expression of any genes encoding the ER enzymes, their product and substrate labeling rates in OE would have increased and decreased, respectively, and *vice versa* in KO. Another possibility is that LHY/CCA1 may activate glycerol-3-phosphate acyltransferase (GPAT) or lysoPA acyltransferase (LPAT) (Figure 3A), which provides the common precursor (LPA or PA) for TAG, DAG, and PC, leading to an increase in all three labeled lipids. However, this can be ruled out by the results from our pulse-chase labeling, where no significant change in any lipid turnover was observed in OE or KO because otherwise, that must be reflected by a difference in the slope of accumulation rates for the affected lipids. Taken together, the metabolic tracking data suggest that FA synthesis is increased in *LHY/CCA1*-OE and decreased in *LHY/CCA1-*KO in plastids.

LHY and CCA1 are generally regarded as transcriptional repressors of their direct targets. However, evidence has also suggested their role in the activation of genes by direct promoter binding. In fact, CCA1 was initially discovered as a transcriptional activator by early studies showing that it bound and activated a light-harvesting chlorophyll *a/b* protein gene (*Lhcb1*3*) in Arabidopsis response to red light (Wang et al., 1997; Green and Tobin, 1999). In addition, the expression of *PRR7* and *PRR9* was dramatically decreased when both *LHY* and *CCA1* were lost but increased by overexpression of either one (Farré et al., 2005). *PRR7* has three CBSs and *PRR9* has an EE, and CCA1 binding to their promoters was verified by EMSA (Farré et al., 2005). The LHY/CCA1 homologs, REVEILLEs (RVE), activate the expression of *PRR5, TOC1*, and *ELF4* by binding to their EE (Farinas and Mas, 2011; Rawat et al., 2011; Hsu and Harmer, 2014). More recent studies also demonstrated the positive effect of LHY/CCA1 on the expression of their direct target genes, including *PHYTOCHROME INTERACTING FACTOR4* (*PIF4*) and *VERNALIZATION INSENSITIVE3* (*VIN3*) (Sun et al., 2019; Kyung et al., 2022).

In addition, the present study suggests that KO and OE of *LHY* and *CCA1* also affect the expression of *BCCP* and *BADC* to impact plastidic ACC that produces the substrate malonyl-CoA for KASIII-catalyzed condensation for *de novo* FA synthesis. A recent RNA-seq study showed that in *lhycca1* the transcript level of *BCCP2* and *KASIII* were decreased with that of *BADCs* slightly up-regulated in Arabidopsis seedlings (Kidokoro et al., 2021), largely in agreement with our transcript analyses in developing seeds. BCCP and BADC are structurally similar and compete for binding to BC, but BADC lacks the biotin-binding site, thus inhibiting the ACC catalytic activity (Salie et al., 2016; Ye et al., 2020). Thus, the opposite effects of LHY/CCA1 on *BCCP2* and *BADC2* transcript levels are consistent with their enhancing role in FA synthesis and TAG production. However, unlike *KASIII*, the promoter regions of *BCCP2* and *BADC2* lack the EE or CBS *cis*-elements, and LHY/CCA1 might regulate the expression of *BCCP2* and *BADC2* via other proteins. CCA1 (and LHY) may form various complexes with other proteins that possibly have differential effects on their target genes, including *BCCP2* and *BADC2*. In fact, many co-regulators, including LIGHT-REGULATED WDs (LWDs), NIGHT LIGHT-INDUCIBLE AND CLOCK-REGULATEDs (LNKs), FAR-RED ELONGATED HYPOCOTYL3 (FYH3), FAR-RED IMPAIRED RESPONSE1 (FAR1), and ELONGATED HYPOCOTYLS5 (HY5), play important roles in the plant molecular clock, and the repressive action of LHY/CCA1 derives in part from their binding to and inhibition of some of the transcriptional activators (Li et al., 2011; McClung, 2019). Moreover, an unknown factor(s), which acts in concert with LHY/CCA1, was proposed to be required for the *VIN3* activation because LHY/CCA1 themselves were not sufficient, and another binding site (‘G-box’) was required for the full activation of *VIN3* (Kyung et al., 2022). Similarly, upon binding to the *PIF4* promoter, CCA1 recruits SHORT HYPOCOTYLS UNDER BLUE1 (SHB1) to mediate the *PIF4* expression. In addition, several Arabidopsis genes are associated with CCA1 (∼400 loci in Col-0) and other sequences in addition to EE/CBS are enriched in them (Nagel et al., 2015; Kamioka et al., 2016). Thus, how LHY/CCA1 affect *BCCP2* and *BADC2* transcript levels remain to be elucidated Previously, we found that PA physically interacted with and inhibited the transcriptional activity of LHY/CCA1 (Kim et al., 2019). The total PA levels in Arabidopsis displayed diurnal fluctuation, and some PA molecular species were under circadian control as their levels did not oscillate in *lhycca1* (Maatta et al., 2012; Kim et al., 2019). Given that PA is a metabolic precursor for TAG biosynthesis, the clock-regulated seed oil production and the interaction between PA and the clock raise the possibility that the circadian clock is under PA-mediated feedback inhibition for fine-tuning TAG production. This is supported by our data showing the inhibitory effect of PA on LHY binding to the *KASIII* promoter (Figure 5E). Conversely, the clock factor-regulated FA synthesis may affect TAG production via PA formation, which also requires FAs. The interaction of PA with LHY/CCA1 and its inhibition on their activation of *KASIII* may serve as a retrograde mediator of the clock regulators to avoid excessive TAG accumulation in response to metabolic and/or environmental cues. Further investigations of the interplays between lipid metabolism and clock regulators have the potential to enhance seed oil production and advance mechanistic understanding of the clock-associated lipid metabolic dysfunctions.

## MATERIALS AND METHODS

### Plant materials and growth conditions

All Arabidopsis plants used in this study were Col-0 ecotype of *Arabidopsis thaliana*. T-DNA insertional double knockout mutant *lhycca1* (*cca1-1 lhy-20*) was generated by and obtained from Pruneda-Paz et al., 2009, and we previously generated the transgenic lines overexpressing *LHY* or *CCA1* (LHY/CCA1-OEs; Kim et al., 2019). All plants were confirmed as homozygotes by verifying no further segregation through PCR-based genotyping and selection of antibiotics (kanamycin or gentamycin)-resistant plants. Seeds were surface-sterilized with 70 % (v/v) ethanol and then with 20 % (v/v) bleach, washed with water, and sown on 1/2 strength of Murashige and Skoog (MS) media supplemented with 1 % (w/v) sucrose and 0.8 % (w/v) agar or on soil (ProMix FPX) filled in 3.25-inch square pots. After stratification at 4 °C for 2 days in the dark, plants were germinated and grown in a growth chamber maintained at 22 °C and a relative humidity of 60% under light cycles of 12-h light/12-h dark with a photosynthetic photon flux density of 120-150 μmol/m^2^/sec.

### Reverse transcription (RT) and quantitative real-time (qRT)-PCR

Total RNA was extracted from plant tissues using TRIzol Reagent (Life Technologies, Carlsbad, CA) according to the manufacturer’s instructions. cDNA was synthesized by High Capacity cDNA Reverse Transcript Kit (Applied Biosystems, Waltham, MA) with 1 μg of RNA and 0.5 μg of oligo(dT)18 primers, according to the manufacturer’s instructions. The reaction was at 37 °C for 2 h with pre-incubation at 25 °C for 10 min and enzyme inactivation at 85 °C for 5 min. The cDNA was amplified with a Taq DNA polymerase using gene-specific primers (sequence provided in Table S1) through the following thermal cycling conditions: pre-incubation at 95 °C for 2 min, 35-40 cycles of 95 °C for 30 sec, 53-60 °C for 30 sec and 68 °C for 1 min, and final extension at 68 °C for 5 min. PCR products were resolved in 1 % (w/v) agarose gel and visualized by staining with ethidium bromide under UV. For quantification, PCR progress was monitored by adding SYBR Green dye using StepOnePlus™ Real-Time PCR System (Applied Biosystems), and data were processed and quantified by StepOne™ Software (v2.0.2). The gene expression was normalized with *ubiquitin10* as an internal standard.

### Nuclear protein extraction

Plant tissue (∼0.5 g) was ground with liquid nitrogen and mixed with 5 mL buffer A (10 mM Tris-HCl pH7.6, 0.5 M sucrose, 1 mM spermidine, 4 mM spermine, 10 mM EDTA, 80 mM KCl). The tissue homogenate was filtered through 4 layers of Miracloth (Calbiochem) and centrifuged at 3,000 ×g for 5 min. After discarding supernatant, the pellet was gently resuspended in 1 mL buffer B (50 mM Tris-HCl pH7.8, 5 mM MgCl_2_, 10 mM β-mercaptoethanol, 20 % (v/v) glycerol). Discontinuous Percoll™ (Amersham Biosciences, Amersham, United Kingdom) gradient was prepared with 2 mL each of 40 % (v/v), 60 %, and 80 % (top to bottom) Percoll™ dissolved in buffer C (25 mM Tris-HCl pH7.5, 0.44 M sucrose, 10 mM MgCl_2_) on 2 mL of 2 M sucrose cushion. The cell suspension was gently loaded on top of the Percoll™ gradient and centrifuged at 4,000 ×g for 30 min. Nuclear layer (light green) right above the 2 M sucrose cushion was carefully taken, washed twice with 1 mL buffer A at 6,000 ×g for 5 min, and then resuspended in 0.2 mL buffer B.

### SDS-PAGE and immunoblotting

Protein samples were mixed with SDS-PAGE loading buffer, boiled for 5 min, and resolved in 10 % (v/v) polyacrylamide gel at 100 V for ∼1 h. Proteins were visualized by staining with Commassie Brilliant Blue followed by destaining with a 3:6:1 (v/v/v) mixture of methanol, water, and acetic acid. For immunoblotting, proteins were electrophoretically transferred onto a polyvinylidene fluoride (PVDF) membrane using Semidry Trans-Blot apparatus (Bio-Rad, Hercules, CA) at 20 V for 20 min. The membrane was blocked in tris-buffered saline with 0.1 % (v/v) Tween-20 (TBST) buffer containing 5 % (w/v) nonfat milk for 1 h, followed by washing three times with TBST buffer. The membrane was incubated for 1 h with primary antibodies: polyclonal anti-LHY (R3095-2, Abiocode, Aguora Hills, CA), polyclonal anti-CCA1 (R1234-3, Abiocode), and polyclonal anti-histone H3 (A01502, GenScript, Piscataway, NJ). After washing three times with TBST buffer, the membrane was incubated with a secondary antibody (anti-rabbit IgG; A7539, Sigma-Aldrich, St. Louis, MO) conjugated with alkaline phosphatase for 1 h. Proteins were visualized by alkaline phosphatase conjugate substrate (Bio-Rad) according to the manufacturer’s instructions.

### Leaf movement measurements

Arabidopsis seedlings were grown on a 1/2 MS plate under T24 cycles for 5 days. They were individually transferred to a 24-well plate by carefully excising an agar square (∼1 cm^2^) surrounding the seedling and transferring it onto the wall of each well, with the cotyledons perpendicular to the plate. In a growth chamber with continuous light, the plate was placed in vertical orientation with a white background to facilitate imaging. Plant images were taken every 90 min for 5 days using Optio-750Z digital camera (Pentax) with the interval shooting mode. Values for vertical coordinates of leaf edge and hypocotyl apex were obtained for each image using the pixel tracking function of Image J software (v1.48), and plotted by Microsoft Office Excel (2010).

### Seed oil content measurements

Approximately 5 mg per sample of mature seeds were accurately weighed and incubated in a glass tube with 2 mL methanol containing 1.5 % (v/v) H_2_SO_4_ and 0.01 % (w/v) butylated hydroxytoluene (BHT) to prevent lipid oxidation at 90 °C for 2 h for transmethylation. Fatty acid methyl esters (FAMEs) were extracted with 1 mL of hexane and 1 mL of water. After a brief centrifugation, the upper phase was transferred into a GC-compatible glass vial and applied to a GC detection system (Shimadzu GC-17A) along with heptadecanoic acid (17:0) as an internal standard for quantification. The GC system was supplied with a hydrogen flame ionization detector and a capillary column SUPELCOWAX-10 (30 m length; 0.25 mm diameter) with helium carrier at a flow rate of 20 mL/min. The oven temperature was maintained at 170 °C for 1 min and then increased in steps to 210 °C, raising the temperature by 3 °C/min. FAMEs from TAG were identified by comparing their retention times with known standards. Each peak area was obtained, converted to lipid amount based on the internal standard amount, and calculated as % (w/w) of the seed weight.

### TAG quantification in developing and germinating seeds

To collect developing seeds, plants grown on soil were marked for flowers when they emerge and siliques were harvested at 5, 7, 9, 11, and 13 DAF, from which 100-150 developing seeds per sample were carefully collected using scalpel and forceps. For germinating seeds, 50-100 seeds or seedlings per sample grown on a 1/2 MS plate were collected at 0, 1, 2, 3, 4, and 5 DAI. The collected tissues were incubated at 75 °C for 15 min in isopropanol containing 0.01 % (w/v) BHT to prevent lipolysis and oxidation. Total lipids were extracted three times with a 2:1 (v/v) mixture of chloroform and methanol with 0.01% (w/v) BHT. All lipid extracts from the same sample were combined and washed with 1 M KCl then with water. The lower organic phase containing lipids were dried under a gentle stream of nitrogen gas and re-dissolved in 0.5 mL of chloroform. TAG was separated from the total lipids by TLC, where 50 μL of the lipid extract was loaded onto a TLC plate (Silica Gel 60 F254, Merck) and developed in a 70:30:1 (v/v/v) mixture of hexane, diethyl ether, and acetic acid until the solvent front reaches near the top of the plate. After drying in a fume hood, the plate was exposed to iodine vapor for visualization of lipids. TAG on the plate was identified based on the position of authentic TAG standard (vegetable oil dissolved in chloroform). TAG spots were scraped out into a glass tube and incubated with a 3:1 (v/v) mixture of methanol and toluene containing 2.5 % (v/v) H_2_SO_4_ and 0.01 % (w/v) BHT at 90 °C for 2 h for transmethylation. FAMEs were then extracted and analyzed by GC as described above.

### Metabolic tracking of radio-labeled lipids

Approximately 100 seeds per sample were collected at 10 DAF as described above and incubated with incubation buffer (IB; 5 mM MES pH 5.7, 1/2 MS medium, 1 % (w/v) sucrose) for 1 h with gentle agitation. 5 μCi (∼86 nmol) of [^14^C]-acetate was added to each sample and further incubated for 0, 0.5, 1, 2, 4, and 6 h, and the seeds were washed with IB three times to remove residual radioisotopes for pulse labeling. For pulse-chase labeling, after incubation with the radioisotopes for 3 h the buffer containing the radioisotopes was discarded, and the seeds were washed with IB three times and further incubated in 1.5 mL of fresh IB for 0, 0.5, 1, 2, 4, and 6 h. The seeds were then mixed with isopropanol and subjected to lipid extraction as described above. Dried lipids were re-dissolved in 0.1 mL of chloroform and separated by TLC as described above with a 70:30:1 (v/v/v) mixture of hexane, diethyl ether, and acetic acid for the separation of TAG and DAG and with a 65:25:4 (v/v/v) mixture of chloroform, methanol, and water for PC. The authentic standards for lipid identification were vegetable oil dissolved in chloroform, 16:0-18:1 DAG, and PC mixture from egg yolk (Avanti Polar Lipids, Birmingham, AL). TAG, DAG and PC spots on the TLC plate were scraped out into a scintillation vial containing 5 mL of scintillation cocktail, and their radioactivity (^14^C) was measured by a scintillation counter (LSC6500; Beckman Coulter, Indianapolis, IN). Chlorophyll contents were calculated as 7.05×A_662_+18.09×A_645_ by measuring the samples spectrophotometrically at 645 nm and 662 nm.

### Chromatin immunoprecipitation (ChIP)

Arabidopsis line with FLAG-tagged CCA1 for ChIP was previously generated (Kim et al., 2019). Approximately 5 g (fresh weight) of 10-day old seedlings grown in 12-h light/12-h dark were harvested at ZT0, frozen in liquid nitrogen, and disrupted using a ball mill (Retsch). Ground tissue was first resuspended in 20 ml crosslinking buffer (60 mM HEPES pH 8.0, 1 M Sucrose, 5 mM KCl, 5 mM MgCl_2_, 5 mM EDTA, 0.6 % (v/v) Triton X-100, 1 mM PMSF, 1 μg/ml Pepstatin, 1 μg/ml leupeptin, 1 μg/ml aprotinin, and 1 mini-Complete protease inhibitor tablet (Roche)), and then after the addition of 1 % (v/v) formaldehyde, was incubated at room temperature for 25 min. Crosslinking was quenched with the addition of 125 mM glycine, pH 8. Crosslinked tissue was filtered through Miracloth and nuclei were pelleted by centrifugation at 4 °C, 3,220 g, for 20 min. Nuclei were resuspended in extraction buffer (0.25 M Sucrose, 10 mM Tris-HCl pH 8.0, 10 mM MgCl_2_, 1 % (v/v) Triton X-100, 1 mM EDTA, 5 mM BME, 0.1-1 mM PMSF, 1 μg/ml Pepstatin, 1 μg/ml leupeptin, 1 μg/ml aprotinin, and 1 mini-Complete protease inhibitor tablet), split into four tubes, and pelleted at 4 °C, 12,000 g, for 10 min. Pelleted nuclei were resuspended in 600 µl SII buffer with protease inhibitors (100 mM NaH_2_PO_4_ pH 8.0, 150 mM NaCl, 5 mM EDTA, 5 mM EGTA, 0.1 % (v/v) Triton X-100, 1 mM PMSF, 1 mini-Complete protease inhibitor tablet, and 50 µM MG132). Nuclei and chromatin preparation proceeded as previously described (Nusinow et al., 2011) with the following modifications. Chromatin was sheared and extracted by sonication (Fisher Scientific) at 50 % power, with 1 sec on/off cycles for 15 seconds (seven cycles for a total of 105 seconds) to obtain an average chromatin size of ∼500 bp. For the immunoprecipitation, samples were incubated with 5 µg anti-FLAG antibody bound to Protein G Dynabeads (Invitrogen).

### Electrophoretic mobility shift assay

To prepare purified LHY protein, *LHY* coding sequence was cloned into pET-53-DEST™ vector (Novagen), expressed in *E. coli* Rosetta™ (DE3), and affinity-purified using Ni-nitrilotriacetic acid (NTA) agarose (QIAGEN) as previously described (Kim et al., 2019). DNA probes used were a 60-bp of *KASIII* promoter region containing each EE or CBS (EE-W1/EE-W2 or CBS-W1/CBS-W2) and the same region with EE/CBS mutated as A↔C and T↔G (EE-M1/EE-M2 or CBS-M1/CBS-M2; full sequence provided in Table S1). Single-stranded complementary oligonucleotides were self-annealed in TAE buffer (10 mM Tris pH 8.0, 50 mM NaCl, 1 mM EDTA) by incubating at 95 °C for 5 min and slowly cooling down to room temperature. Ten picomole of the resulting double-stranded DNA was incubated in TAE buffer with 1-20 μg of the purified LHY and 0.1-10 μg of sonicated lipids (PA mixture from egg yolk, 18:1-18:1 PA, or 16:0-18:1 PA) at room temperature for 15 min. The reaction mixture was mixed with gel loading buffer (30 % (v/v) glycerol without bromophenol blue), and DNA was separated in 1 % (w/v) agarose gel at 100 V at 4 °C and visualized by staining with ethidium bromide under UV light.

### Transactivation assay

For effectors, *LHY* coding sequence was cloned into p35S-FAST-GFP as described previously (Kim et al., 2019), and the empty vector was used as a control. For reporters, promoter region of *KASIII* (1,774 bps upstream from start codon) was PCR-amplified from Arabidopsis genomic DNA and cloned into pENTR-MCS at *NotI* and *AscI* sites and then into pFLASH-Fixed containing *LUCIFERASE* by Gateway™ cloning using LR Clonase™ (Life Technologies). TOC1::LUC was generated previously (Kim et al., 2019). The DNA constructs were transformed into Agrobacteria (*Agrobacterium tumefaciens* C58C1) by cool shock in liquid nitrogen. The transformed cells were suspended in infiltration buffer (10 mM MgSO_4_, 10 mM MES pH5.2) containing 100 μM acetosyringone and infiltrated into ∼3-week old tobacco (*Nicotiana benthamiana*) leaves using a 1-ml syringe without needle. After two days the infiltrated regions on the leaves were excised, from which total proteins were extracted and measured for concentration by Bradford assay. 50 μg of the proteins was mixed with 1 mM luciferin in a 96-well plate with dark bottom and measured for luminescence intensity using a microplate reader (Infinite M200pro, Tecan). Fluorescence intensity was measured by SpectraMax® M2 (Molecular Devices) with excitation at 485 nm and emission at 538 nm.

### Accession numbers

Sequence data from this article can be found in The Arabidopsis Information Resource (http://www.arabidopsis.org) under accession numbers: LHY, At1g01060; CCA1, At2g46830; TOC1, At5g61380; ACC1, At1g36160; ACC2, At1g36180; KASI, At5g46290; KASII, At1g74960; KASIII, At1g62640; BC, At5g35360; BCCP1, At5g16390; BCCP2, At5g15530; BADC1, At3g56130; BADC2, At1g52670; BADC3, At3g15690; α-CT, At2g38040; β-CT, Atcg00500; Ubiquitin10, At4g05320

## SUPPLEMENTAL DATA

**Figure S1.**
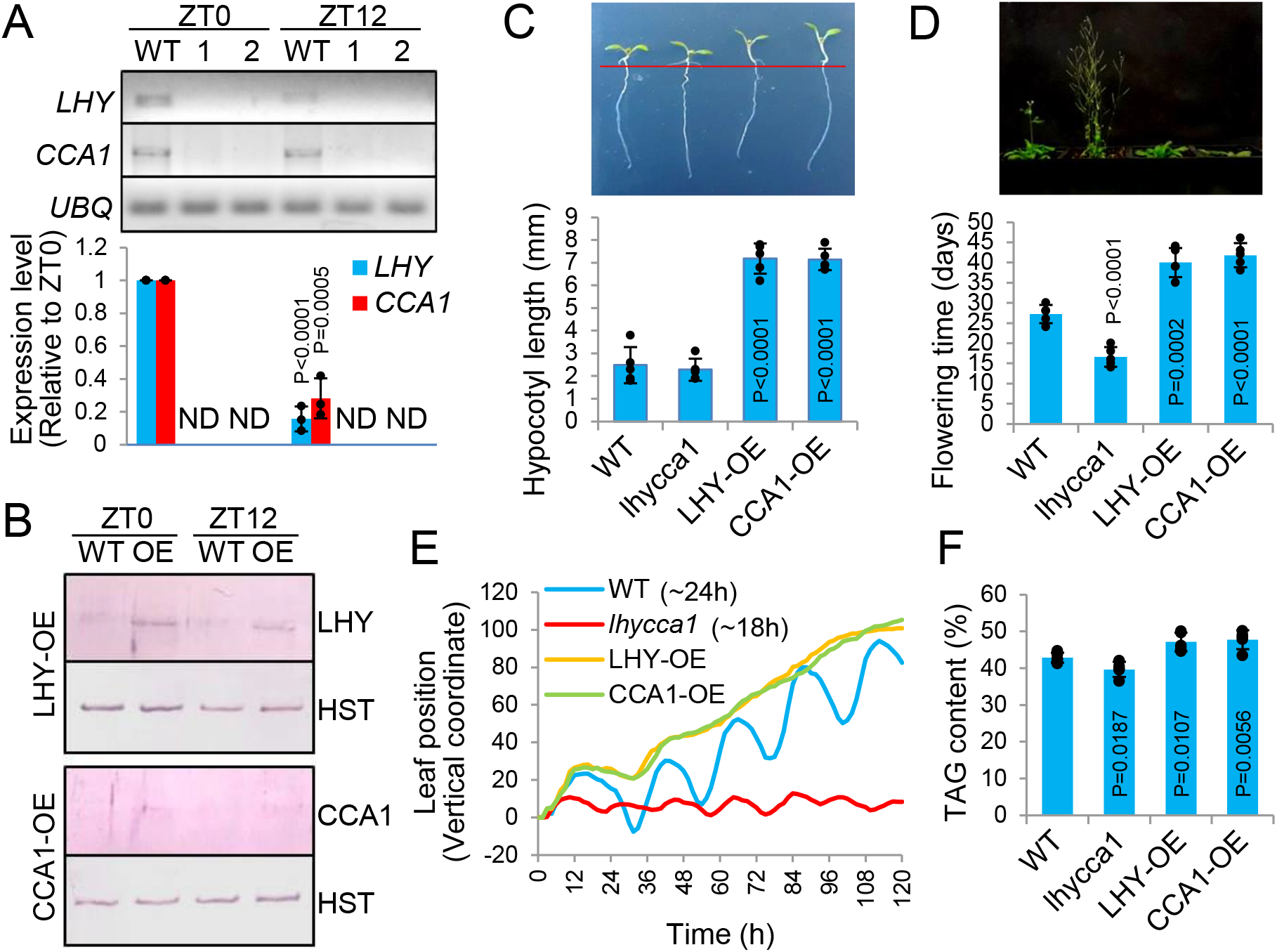
Confirmation of the *LHY*/*CCA1*-altered plant materials. **A**. PCR confirmation of *lhycca1*. Total RNA was extracted from 7-day old WT and two independent plants of *lhycca1* (1 and 2) at the indicated Zeitgeber time (ZT). Reverse transcription PCR (top) and quantitative real-time PCR (bottom) were performed using gene-specific primers. *Ubiquitin10* (*UBQ*) was used as a loading control (top) and internal standard (bottom). Values are mean ± SD with individual data points and *p* values denoting statistical significance (to ZT0; n=3). ND; not detected. **B**. Immunoblotting to confirm LHY/CCA1-OEs. Total nuclear proteins were extracted from 7-day old WT and LHY/CCA1-OEs at the indicated ZT. Immunoblotting was performed using protein-specific antibodies. Histone H3 (HST) was used as a nuclear marker. **C** & **D**. Phenotypic analyses of *LHY*/*CCA1*-altered plants. Hypocotyl length of 7-day old seedlings (C) and time taken to first flower (D) were measured for WT, *lhycca1*, and LHY/CCA1-OEs. Values are mean ± SD with individual data points and *p* values denoting statistical significance (to WT; n=5). Images are 7-day (C) and 25-day (D) old Arabidopsis plants.**E**. Vertical leaf movement. Plants were entrained to 12-h light/12-h dark cycle for 5 days. Leaf movement was monitored under constant light and is shown here as normalized to initial leaf position. Period length is in parenthesis. Data for WT and *lhycca1* are reproduced from Kim et al., 2019. **F**. Seed oil contents of dry seeds from the indicated plant lines measured by nuclear magnetic resonance. Values are mean ± SD with individual data points and *p* values denoting statistical significance (to WT; n=5).

**Figure S2.**
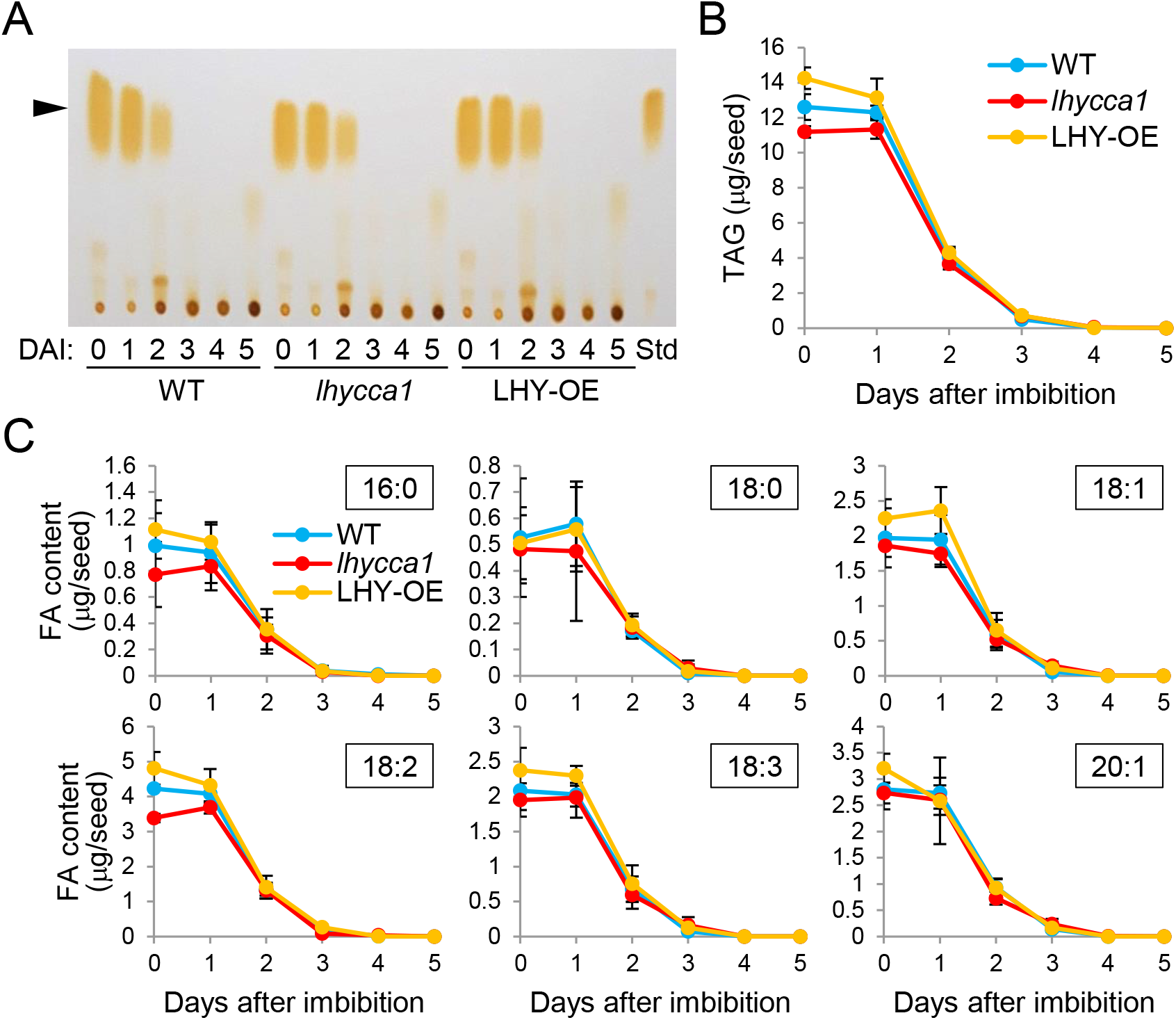
TAG degradation during seed germination. **A**. Thin layer chromatography (TLC) separation of TAG. Total lipids were extracted from germinating seeds of WT, *lhycca1*, and LHY-OE at the indicated days after imbibition (DAI). The lipids were separated by TLC and visualized by iodine exposure. Arrow head indicates the position of TAG. Std, authentic TAG standard. **B**. Contents of total TAG. TAG spots in A were scraped out and quantified by gas chromatography for TAG contents. Values are mean ± SD (n=3). **C**. Contents of individual fatty acids. The data in B are shown here as the amounts of individual species of fatty acids (number of carbons : number of double bonds). Values are mean ± SD (n=3).

**Figure S3.**
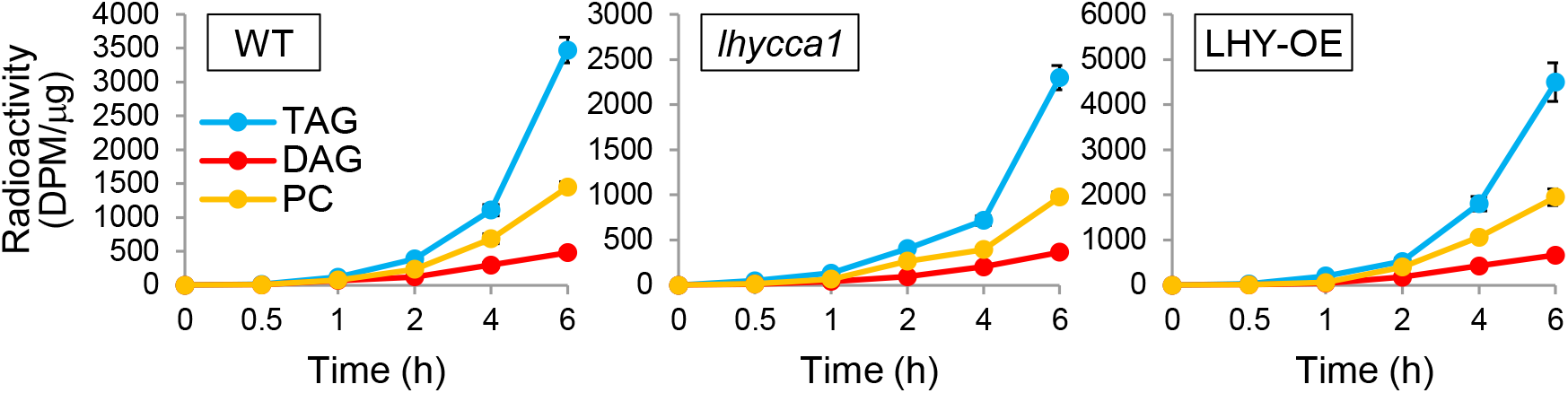
Relative labeling rates of TAG, DAG, and PC in *LHY*/*CCA1*-altered plants. Graphs in Figure 3B have been modified here to show TAG, DAG, and PC on a single plot for each of WT, *lhycca1*, and LHY-OE.

**Figure S4.**
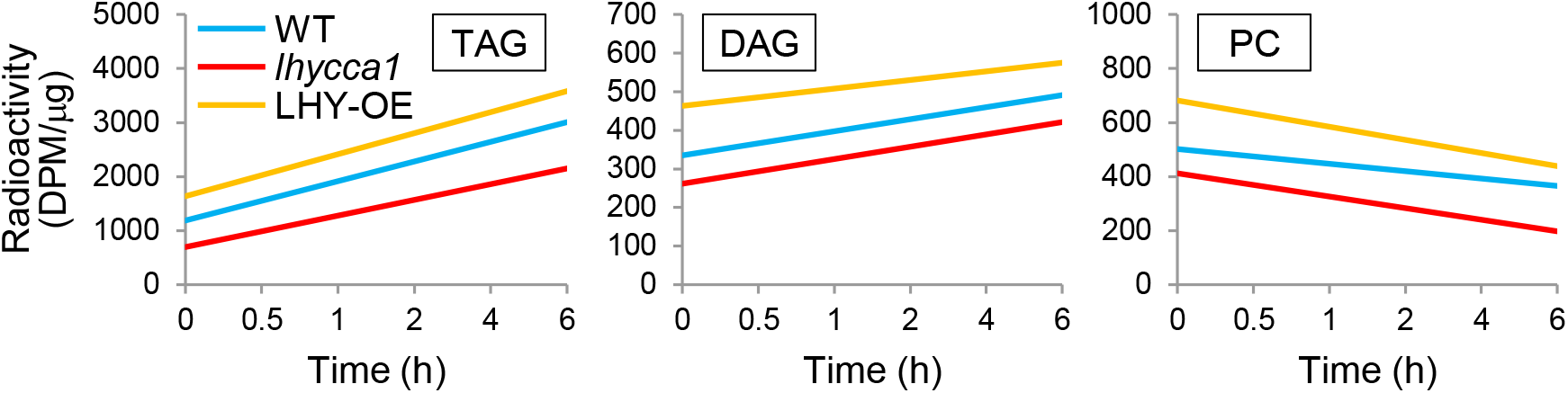
Trend lines of radioactive lipid formation in pulse-chase labeling. Graphs in Figure 3C have been modified here to show them as trend lines over time.

**Figure S5.**
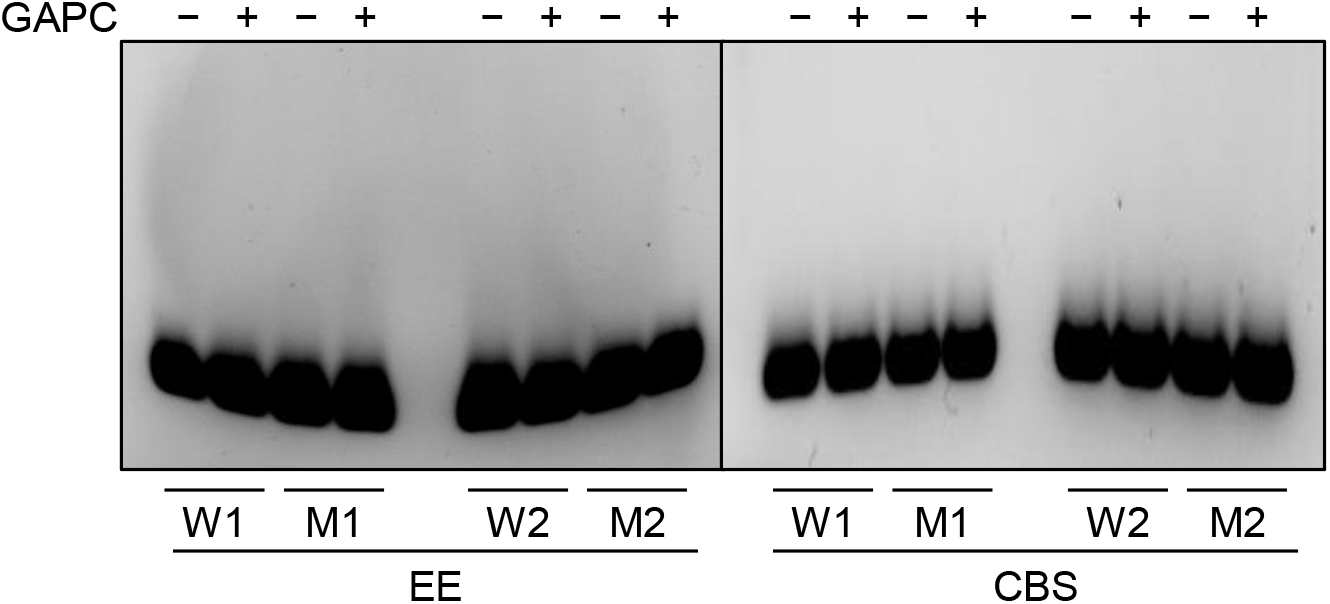
Negative controls for EMSA. 60-bp DNA fragments containing wild-type (W) and mutated (M) EE and CBS were incubated without (-) and with (+) 20 μg of purified glyceraldehyde-3-phosphate dehydrogenase (GAPC). The DNA was separated in an agarose gel and visualized by ethidium bromide. 1 and 2 indicate the first and second EE or CBS, respectively, as shown in Figure 5A.

**Table S1.**
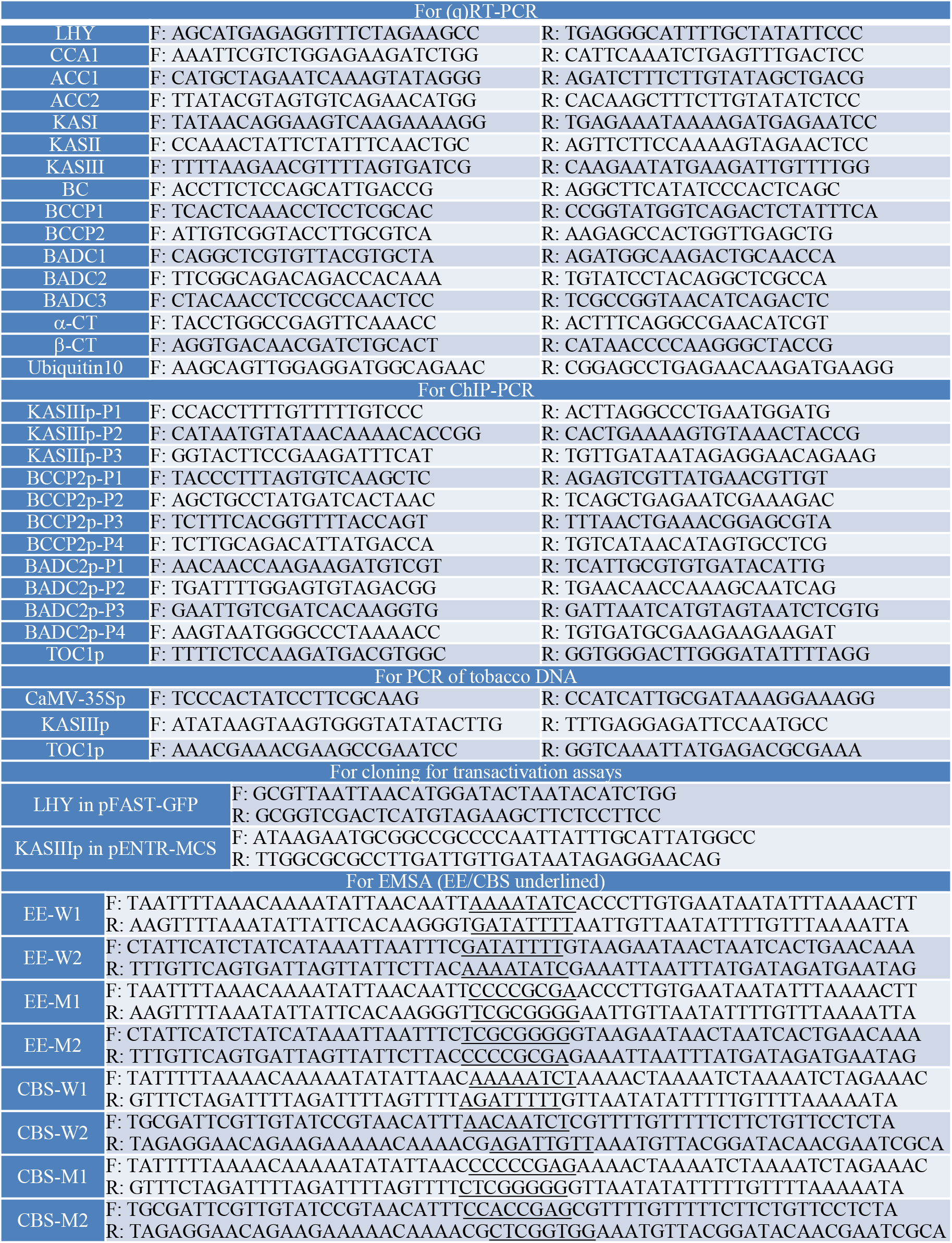
Sequence of oligonucleotides used in this study (5’→3’)

## ACKNOWLEDGMENTS

We thank Maria L. Sorkin for providing plant materials for and assisting with chromatin immunoprecipitation. Research reported in this article was supported by the National Institute of

General Medical Sciences of the National Institutes of Health under award number R01GM141374.

## AUTHOR CONTRIBUTIONS

S.K. designed and performed research, analyzed data, and wrote the paper; D.A.N. analyzed gene expression data, discussed data and ideas, and edited the manuscript; K.N.E. performed chromatin immunoprecipitation and edited the manuscript; and X.W. proposed and supervised research and edited the paper.

The authors declare no competing interest.

